# Microbial diversity and insights into feeding behavior of the two-spot cotton leaf hopper (*Amrasca biguttula*)

**DOI:** 10.64898/2026.07.28.741253

**Authors:** Saptarshi Ghosh, Vamsidhar Reddy Netla, Udaya Durga Prasad Kodigepalli Suresh, Kamran Rashid, Gulshan Kumar, Jeffrey A. Davis, Rajagopalbabu Srinivasan

## Abstract

The recent invasion of the two-spot cotton leaf hopper (TSCL) (*Amrasca biguttula*) into the U.S. now threatens row and vegetable crops as well as ornamentals. TSCL ecology and biology, including microbial diversity, are understudied. Insect-specific microbes could influence host biology and ecology including insect-plant interactions. Here, microbiota (bacteria and viruses) within TSCL along with its feeding behavior were characterized using high throughput sequencing, microscopy, and Electrical penetration graph (EPG). Results revealed low diversity of bacteria dominated by *Wolbachia* and confirm the absence of obligate (primary) symbionts. Two distinct and differentially abundant *Wolbachia* strains (B-supergroup) were found co-infecting midgut tissues of 60% of TSCL tested, and the remaining 40% were free of symbionts. Virome analysis uncovered a diverse complex of at least eight dsDNA genomes (12-17 kb) of Adintoviruses (*Eupolintoviridae*), but no RNA viruses commonly found in other leafhoppers were identified in TSCL. The mesophyll feeding nature of TSCL was identified by observing chlorophyll derived autofluorescence in midguts and higher abundance of chloroplast transcripts (photosystem I, II, Rubisco, cytochrome b6 and ATP synthase complex) in TSCL (whole insects) compared with thrips and aphids. EPG recordings show a cell-rupture style of feeding with differences between adults and nymphs. Lastly, consistent with its cell-rupturing feeding style, no evidence for the presence of plant-pathogenic bacteria or viruses that could induce hopperburn was found. These results overall provide insights into the microbial diversity and feeding pattern of TSCL and lay a foundation for future research.

**Key message:** This work characterizes the TSCL microbiome and its mesophyll feeding pattern. Symbionts within insects often complement the nutritional deficiencies of its host and can aid insect spread by boosting immunity. The findings of this study lay the foundation for future research on the biology, ecology, and management of this invasive pest.

## Introduction

*Amrasca biguttula* (Ishida), the two-spot cotton leafhopper (TSCL) (Cicadellidae, Auchenorhynca, Hemiptera), is a polyphagous sap sucking pest that causes severe economic losses to wide range of crops including cotton (1), vegetables (2, 3) and ornamentals (4, 5). Originally native to Asia, TSCL was first detected in the USA in 2023 in Puerto Rico (6). By July 2025, it had spread to the majority of Southern States, ranging from Texas to Virginia (7). Based on sequencing data, it appears that a single haplotype of TSCL is responsible for the invasion, and the same haplotype is also the most widespread globally (8). Factors driving the spread of this invasive haplotype in the USA are yet to be understood.

Hemipteran insects rely on obligate (primary) endosymbiotic bacteria to complement their nutritionally deficient phloem/xylem diets (9, 10). These symbionts compensate for nutritional imbalances and allow for diversification of insect species by facilitating or restricting access to new feeding niches (11). Cicadellidae members commonly harbor a primary bacterial symbiont, *Candidatus* Sulcia muelleri along with a companion symbiont *Candidatus* Nasuia deltacephalinicola in specialized cells called bacteriocytes (12, 13). Presence of *Sulcia*, a strict requirement in xylem feeders, is lost and replaced with other bacterial or fungal symbionts in cicadellids that have transitioned to phloem and/or parenchyma diets (14).

Other than obligate symbionts, cicadellids also harbor secondary symbiotic/facultative bacteria (15, 16) in bacteriocytes or other tissue such as the fat bodies, midgut, or reproductive organs (17, 18). The role of secondary symbionts in Cicadellids is yet to be fully understood; however, in other hemipterans they confer fitness benefits (19), aiding in defense against abiotic (20) and biotic stress (21–23). In contrast, they also can confer negative impacts on host biology by competing for resources, resulting in reduced adult emergence and slower development (24, 25). Secondary endosymbionts are primarily maintained through vertical transmission, but evidence of horizontal spread also exists (26–28). Other than direct influence on insect nutrition, abiotic/biotic defense and developmental processes, bacterial symbionts can indirectly influence insect invasiveness by broadening their host plant range, mitigating plant defense and enhancing insecticide resistance (29–31).

The two-spot cotton leafhopper (TSCL) belongs to *Typhlocybinae* –the second-largest subfamily within Cicadellidae (32), which differ from other leaf hoppers due to their use of mesophyll as the primary feeding niche instead of the vascular tissue (32–34). Absence of any primary bacterial symbiont differentiates *Typhlocybinae* members from other leaf hoppers, and the absence is in part attributed to their nutrient rich mesophyll diet (17, 35). Potato leaf hopper (*Empoasca fabae*)is the only known *Typhlocybinae* exception that harbors *Sulcia* –the primary symbiont (36).

Characterization of microbial diversity within TSCL and its feeding biology is necessary to reaffirm the relationship between nutrition and symbionts. Moreover, elucidation of microbial diversity within TSCL is important for understanding factors driving its spread across continents and haplotype divergence. Thus, this study assessed the microbiota (bacteria fungi and viruses) within TSCL by high throughput sequencing. In addition, to understand the injury potential of TSCL, probing and feeding patterns were characterized using high throughput sequencing, microscopy, and Electrical penetration graph (EPG). Lastly, since many leafhoppers transmit plant pathogens, this study also examined the presence of TSCL transmitted plant pathogenic bacteria and/or viruses.

## Materials and Methods

### Collection and maintenance of TSCL colony

A laboratory population of *A. biguttula* was maintained for a minimum of four generations since its inception from field collection of adults (N∼25) in September 2025 from cotton and eggplant at UGA Plant Sciences farm, Tifton, GA. The TSCL colony was maintained on cotton plants cv. PHY 339 WRF (Corteva, Indianapolis, IN, USA) in insect-proof cages (47.5 x 47.5 x 93 cm^3^ BugDorm-4E4590 insect rearing cages, Megaview Science Co. Taichung, Taiwan) in greenhouses at the University of Georgia, Griffin, GA at 27 ± 5 °C, 60% RH, and 14 h L:10 h D photoperiod.

### Microbiome analyses

#### Bacteria diversity

High throughput amplicon sequencing of the variable region (V3-V4 region) and full length 16S rDNA was performed for identification of bacteria within TSCL. Identified sequence variants of bacteria were verified by restriction fragment length polymorphism (RFLP) of PCR amplified full length 16S rDNA using generic primers. Bacteria diversity within symptomatic TSCL-infested plant samples also was investigated by amplicon sequencing of 16S rDNA (V3-V4) to identify the presence of any TSCL transmitted plant pathogenic bacteria/phytoplasmas.

### Amplicon sequencing of V3-V4 region of 16S rDNA

V3-V4 region (450 bp) of 16S rDNA were amplicon sequenced (250 bp paired end reads) from genomic DNA samples (N=2) extracted from the laboratory TSCL (10 adults each) population reared on cotton. V3-V4 region was PCR amplified using barcode-tagged primers (Table S1) and sequenced on Illumina Novaseq 6000 sequencing platform (Illumina, San Diego, CA, USA) following the metagenomic amplicon sequencing service pipeline at Novogene Corporation Inc. (Sacramento, CA).

Bioinformatic analyses were carried out using QIIME2 v2025.4 (quantitative insights into microbial ecology) platform (37). The adapters and primers were removed from paired end reads using QIIME2 plug in q2-cutadapt (37). The cleaned reads were quality filtered, dereplicated, and chimeras were removed and merged using QIIME2 plug in dada2 (qiime dada2 denoise-paired) to obtain amplicon sequence variants (ASVs). The ASVs were classified using the Qiime2 Naïve Bayesian classifier against silva 16s/18s SSU database (silva release 138.2). ASVs matching to chloroplast and mitochondria were not considered for downstream analysis. Identified ASVs from TSCL were compared with publicly available data of other *Typhlocybinae* members (*Fagocyba cruenta*, *Kybos virgator, Linnavuoriana sexmaculata, Notus flavipennis, Zygina hyperici* and *Zygina rubrovittata*) based on minimal number of ASVs (79,907) and presence of *Wolbachia* (17). The ASVs of TSCL and other *Typhlocybinae* members were rarefied to sample depth of 79,907 and used for beta diversity Bray-Curtis dissimilarity analysis. Beta diversity analyses with Bray-Curtis dissimilarities were performed using QIIME2 diversity plug in core-metrics-phylogenetics. Principal coordinate analysis (PCoA) plot with beta diversity Bray-Curtis distances plots were generated using R packages and R scripts (38). The statistical difference of beta diversity was performed with permutational multivariate analysis of variance (PERMANOVA) with Mann-Whitney U test.

Bacteria diversity within field collected leaf samples showing puckering and distortion symptoms from cotton (N=4), eggplant (N=2) and okra (N=2) plants infested with TSCL also were analyzed using V3-V4 amplicon sequencing pipeline. Leaves from respective plant species without insect infestation were used as control.

#### Amplicon sequencing of full length 16S rDNA

Genomic DNA samples (N=2) extracted from TSCL (10 adults each sample) were used as template for amplicon sequencing of complete 16S rDNA using amplicon metagenomic sequencing services at Novogene Corporation Inc. (Sacramento, CA). Near full-length 16S rDNA was PCR amplified using barcoded adapter primers (Table S1), purified, pooled in equimolar amounts prior to library preparation using Kinnex long-read sequencing technology. Libraries meeting quality standards were sequenced on the PacBio Revio platform using Single Molecule, Real-Time (SMRT) sequencing to obtain high-quality long-read data.

Raw PacBio HiFi reads were processed using the DADA2 plugin within the QIIME2 pipeline (qiime dada2 denoise-ccs). This workflow included denoising, which encompassed quality filtering, dereplication, and chimera removal to identify amplicon sequence variants (ASVs). Taxonomic assignment of the identified ASVs was deduced using a Naïve Bayesian classifier trained against SILVA 138.2 (16S/18S) rDNA database. The classified ASVs and resulting feature tables were subsequently utilized for further downstream diversity and phylogenetic analyses.

#### Validation of *Wolbachia* isolates by PCR-RFLP of full length 16S rDNA and partial Wolbachia surface protein (Wsp) gene

DNA extracted from pooled samples (N=10) was used as template to PCR amplify a 1.5 kb fragment of eubacterial 16S rDNA using universal primers (39) and partial Wsp *Wolbachia* gene fragment using 81F and 471R primers (Table S1). The PCR fragments were ligated to pJET1.2/blunt cloning vector (ThermoFisher Scientific), transformed to *E. coli,* and positive bacterial colonies with inserts were confirmed by PCR using pJET1.2 forward and reverse sequencing primers. Positive colonies for 16S rDNA (N=70) and Wsp (N=40) were screened by RFLP using *Taq*I and *Alu*I restriction enzymes, respectively, as previously described (39). Plasmids extracted from a minimum of three clones showing unique RFLP profiles were Sanger sequenced in both directions using pJET forward and reverse sequencing primers and analyzed for nucleotide sequence differences.

#### Amplicon sequencing of full length fungal ITS

Fungal diversity within TSCL samples was captured by amplicon sequencing of the complete ITS region using amplicon metagenomic sequencing services at Novogene Corporation Inc. (Sacramento, CA). Universal primers (Table S1) with barcoded adapter primers were used to PCR amplify complete ITS and sequenced on the PacBio Revio platform using Single Molecule, Real-Time (SMRT) sequencing. Raw PacBio HiFi reads were processed using the DADA2 plugin within the QIIME2 pipeline (qiime dada2 denoise-ccs) as described above.

#### PCR based detection of *Wolbachia,* quantitation of sequence variants and phylogeny

To determine the frequency of TSCL harboring *Wolbachia*, DNA extracts from individual adults (40) were screened for presence of *Wolbachia* by PCR using 16S rDNA specific primers (Table S1). *Wolbachia* sequence variants, WSV1 and WV2 were specifically detected and quantitated within individual TSCL samples by qPCR using specific primers (Table S1) that amplify Wsp gene fragment of the WSV1 and WSV2. Relative quantities of WSV1 and WSV2 normalized to the ribosomal protein S13 (RPS13) gene (Table S1) were calculated using ΔΔct method, and significant differences were inferred using Wilcoxon’s rank sum test.

Nucleotide sequences of 16S rDNA, *wsp* and *ftsz* gene of *Wolbachia* obtained from laboratory TSCL, were aligned with sequences from other arthropods available in GenBank. The best-fit nucleotide substitution model was identified according to Bayesian Information Criterion score (BIC), and phylogenetic trees were constructed using maximum likelihood with 1000 bootstrap replicates using IQ-TREE v3.1.0 (41). The resulting consensus trees were visualized using iTOL v7 (Interactive Tree of Life) (42).

#### Localization of *Wolbachia* within TSCL and midgut tissues

*Wolbachia* within whole TSCL adults were localized by fluorescent in situ hybridization (FISH) using a cyanine 3 labeled *Wolbachia* specific DNA probe (Table S1) or without probe (control) as described in (43). Briefly, whole insects were fixed in Carnoy’s fixative (chloroform: ethanol: glacial acetic acid, 6:3:1, v/v) overnight, cleared with 6% hydrogen peroxide in ethanol, and incubated at room temperature with 100 nM of probe in hybridization buffer (20 mM Tris pH 8, 0.9 M NaCl, 0.01% SDS, 30% formamide). Samples were mounted on slides with hybridization buffer containing 100 μg/ml DAPI and viewed under Zeiss LSM700 confocal laser scanning microscope. *Wolbachia* was immunolocalized within fixed (4% paraformaldehyde) TSCL midgut tissues using 2 μg/ml of *Wolbachia* FtsZ polyclonal antibody (ThermoFisher Scientific) and 4 μg/ml of Alexa Fluor^TM^ 488 conjugated goat-anti-rabbit secondary antibody as described earlier (44). Midgut samples stained only with the secondary antibody were used as a control. Mounted midgut samples were viewed using a Zeiss LSM700 confocal laser scanning microscope.

### RNA sequencing of TSCL and host plants for RNA virus discovery

Leaf samples from field grown (UGA plant sciences farm, Tifton, GA) cotton, eggplant and okra infested with TSCL showing puckering and distortion symptoms, and laboratory TSCL population and cotton plants used for rearing them, were collected and subjected to virome analysis for identification of prospective TSCL-transmitted plant viruses. Total RNA extracted from TSCL using RNeasy micro kit (Qiagen, GmbH, Hilden, Germany) and from plant samples using Spectrum Plant Total RNA Kit (Sigma-Aldrich Inc. St. Louis, MO, USA). Extracted RNA was treated with Ribo-zero rRNA removal kit (Illumina, San Diego, CA, USA) for ribosomal RNA (rRNA) depletion and used for library preparation for Illumina deep sequencing (Novogene Corporation Inc. Sacramento, CA, USA). cDNA libraries were prepared using random hexamers and paired end (150 bp) sequenced on Illumina Novaseq X Plus platform (Illumina, San Diego, CA, USA).

Raw reads were quality controlled using FastQC v0.12.0 (45) and MultiQC v1.28 (46), adapters were removed and trimmed using fastp v0.23.4. Processed reads were assembled into contigs using either reference-based or *de novo* assembly. Virus reads were enriched by removal of host reads by mapping total reads to respective reference genomes. Reads from cotton and okra were mapped to their available respective reference genomes (cotton: GCF_007990345.1, okra: GCA_035048815.1) from NCBI. Unmapped reads were extracted using bowtie2 v2.5.4 with default parameters and assembled into contigs using *de novo* assembler SPAdes v4.1.0 (47) with the rnaviralSPAdes parameter and K-mers of 21,33,55 and 77.

For TSCL and eggplant, total trimmed reads were *de novo* assembled into contigs using SPAdes v4.1.0 (47) with the rnaviralSPAdes parameter and K-mers of 21,33,55 and77 due to the unavailability of a reference genome. Assembled contigs were analyzed by BLASTX algorithm search against non-redundant (nr) virus protein databases in DIAMOND v2.1.15 (48). Open reading frames (ORFs) were identified from assembled virus contigs using the NCBI ORF finder algorithm. Translated proteins from the ORFs were compared with other virus proteins in the non-redundant protein database by BLASTP and to predicted virus domains using NCBI conserved domain database (CDD) and HHpred search. The virus contigs were mapped back to raw reads using Bowtie2 v2.5.4 to check virus read abundance and depth.

### Comparison of autofluorescence from chloroplast pigments in midguts

To assess mesophyll tissue feeding, TSCL adults reared on cotton plants were separately fed on either artificial diet (15% sucrose) for 48 hours to clear guts or attached to an excised cotton leaf with petiole dipped in water with a clip cage for 48 hours. Next, midguts from diet or leaf-fed insects were extracted in 1X PBS and then fixed for 45 minutes with 4% paraformaldehyde, washed thrice with PBST (1XPBS + 0.05% Tween 20) and mounted with PBS containing DAPI 100 μg/ml on glass slides with coverslips. Leaf and diet fed midguts were visualized under Fein-Optic FZ8 stereo-trinocular microscope and imaged to compare chlorophyll derived green pigmentation. Confirmation of green autofluorescence was conducted using a Zeiss LSM700 confocal laser scanning microscope using 555 nm excitation and 583 nm emission.

### Comparison of chloroplast transcripts in RNA-seq libraries of TSCL with other phloem feeders

Abundance of chloroplast specific transcripts of TSCL with other phloem feeding insect pests of cotton were compared to confirm mesophyll feeding by TSCL. The RNA-seq reads of TSCL generated in this study and RNA-seq datasets from cotton-reared western flower thrips, *Frankliniella occidentalis* (epidermal feeder) (unpublished) and cotton aphid, *Aphis gossypii* (phloem feeder) (49) generated previously, were compared for abundance of chloroplast specific transcripts. Processed reads were mapped to the cotton reference genome (*Gossypium hirsutum* L.: GCF_007990345.1) using splice aware STAR RNA-seq aligner (50). Mapped reads were assembled and annotated using a reference-guided approach in StringTie (51) by utilizing the cotton reference genome as a scaffold. The resulting transcripts were compared and consolidated against the reference annotation using Gffcompare to ensure structural accuracy. Transcript abundance for chloroplast-related genes was subsequently quantitated using StringTie, with expression levels normalized as Transcripts Per Million (TPM) to facilitate downstream comparative analysis.

### Electrical penetration graph (EPG) recordings

TSCL adults (1-10 days old) and nymphs (4-5^th^ instar) were attached to 3.5 cm long, 12 μM diameter gold wire (0.0005-inch diameter, Sigmund Cohn Co. Mount Vernon, NY, USA) using hand formulated water-based silver glue (https://www.epgsystems.eu/files/SILVER%20GLUE%20recipe.pdf) following protocols as outlined in Walker et. al 2022 (52). Post wiring, insects were starved for one hour and placed on abaxial surface of the top leaf of cotton cv. PHY 339 WRF (4-5 leaf stage), and EPG waveforms were recorded for 10 hours using a 4-channel AC/DC EPG monitor (EPG Technologies, Gainesville, FL, USA), one TSCL per channel, and DI-710 USB data acquisition system (Dataq Instruments, Akron, OH, USA) at 25-30 mV DC substate voltage, 75X gain and 10^9^ Ω input resistance.

## Results

### Bacteria diversity in TSCL

Good’s coverage (>99.99%) estimates of high-throughput sequencing of the 16S rDNA gene indicated that the sampling depth sufficiently captured total bacteria diversity and revealed that a low-diversity bacteria community overwhelmingly dominated by *Wolbachia* reside within TSCL. In both the V3-V4 dataset (254,802 amplicon sequences) and the full-length PacBio dataset (9,166 amplicon sequences), most reads belonged to *Wolbachia* (Fig. 1A, 1B; Table S2). Notably, no obligate bacterial symbiont (*Baumannia*, *Sulcia*, *Zinderia*, *Nasuia*) or other facultative endosymbionts commonly associated with *Cicadellidae* insects could be detected. Bacterial diversity structure compared using PERMANOVA analysis of 16S rDNA sequences showed no significant differences between TSCL and other *Typhlocybinae* members, with TSCL sharing the closest taxonomic abundance with *Kybos virgator*, a leafhopper infesting willow trees in Poland (Fig. 1C). PCR screening of individual TSCL samples with *Wolbachia* specific 16S rDNA confirmed that only 60.0% (18/30) of individuals harbor *Wolbachia* (Fig. 1D), while the rest were devoid of secondary symbionts.

**Fig. 1:**
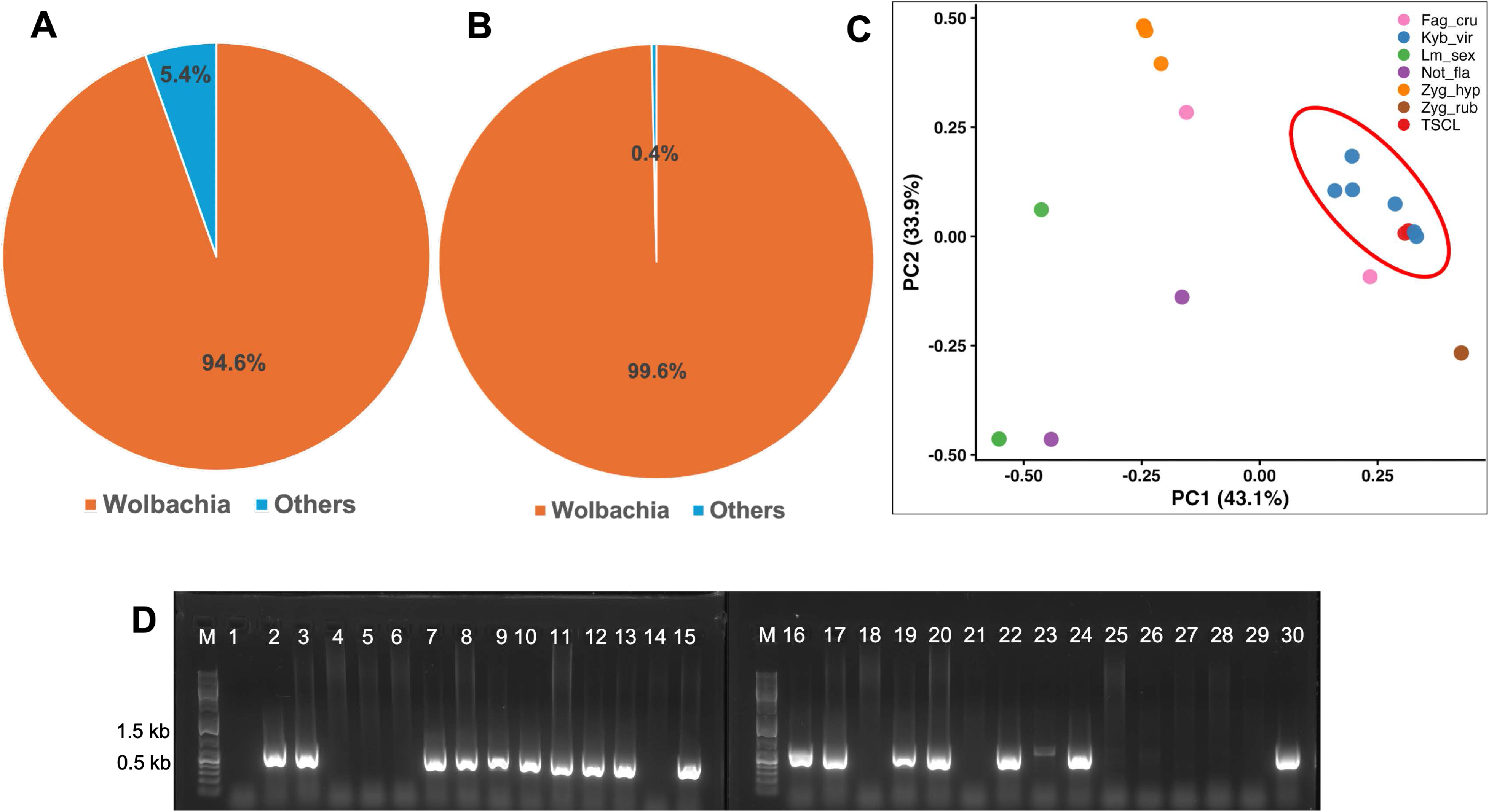
Percentage composition of bacterial genera captured by V3-V4 segments (A) and full length (B) 16S rDNA amplicon sequencing. Principal co-ordinate analysis (PCoA) of Bray-Curtis dissimilarities showing bacterial community compositions of TSLH and other *Wolbachia* harboring *Typhlocybinae* members (C). PCR screening for *Wolbachia* prevalence in individual TSCL adult samples (N=30) using 16S rDNA specific primers (D).

Full-length 16S sequencing identified two *Wolbachia* sequence variants, designated as WSV1 and WSV2 in this study, with WSV1 being the dominant variant (71% of *Wolbachia* reads) (Fig. 2A, Table S2C). Presence of the two *Wolbachia* isolates was validated by PCR-RFLP of cloned full length 16S rDNA and Wsp partial gene fragment, which yielded unique profiles for the more abundant WSV1 and the less abundant WSV2 isolates. Notably, qPCR analysis using variant-specific primers targeting the Wsp gene confirmed that both isolates are co-harbored within TSCL individuals, albeit WSV1 was twice as abundant as WSV2. Phylogenetic analysis of *Wolbachia* 16S rDNA, Wsp and FtsZ nucleotide sequences generated in this study placed both WSV1 and WSV2 within the ‘B’ subgroup of *Wolbachia* (Fig. 3A, B, Fig. S1). The less abundant WSV2 isolate clustered closely with *Wolbachia* isolates previously identified from TSCL in Pakistan, whereas the dominant WSV1 formed a distinct clade (Fig. 3 A, B, S1).

**Fig. 2:**
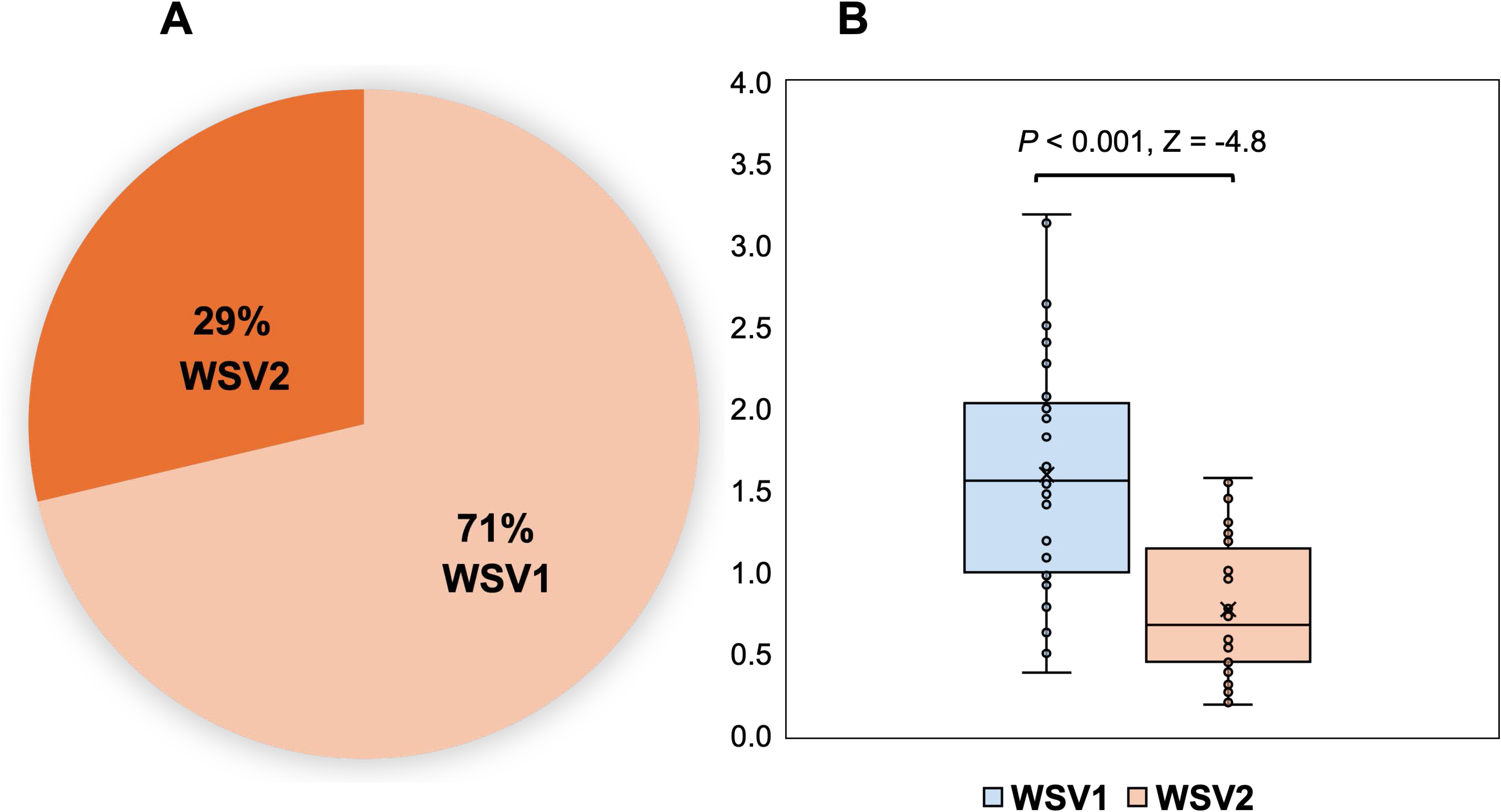
Percentage abundance of *Wolbachia* sequence variants identified by full length 16S rDNA amplicon sequencing (A) and relative quantitation of variants normalized to rps13 gene by qPCR (B).

**Fig. 3:**
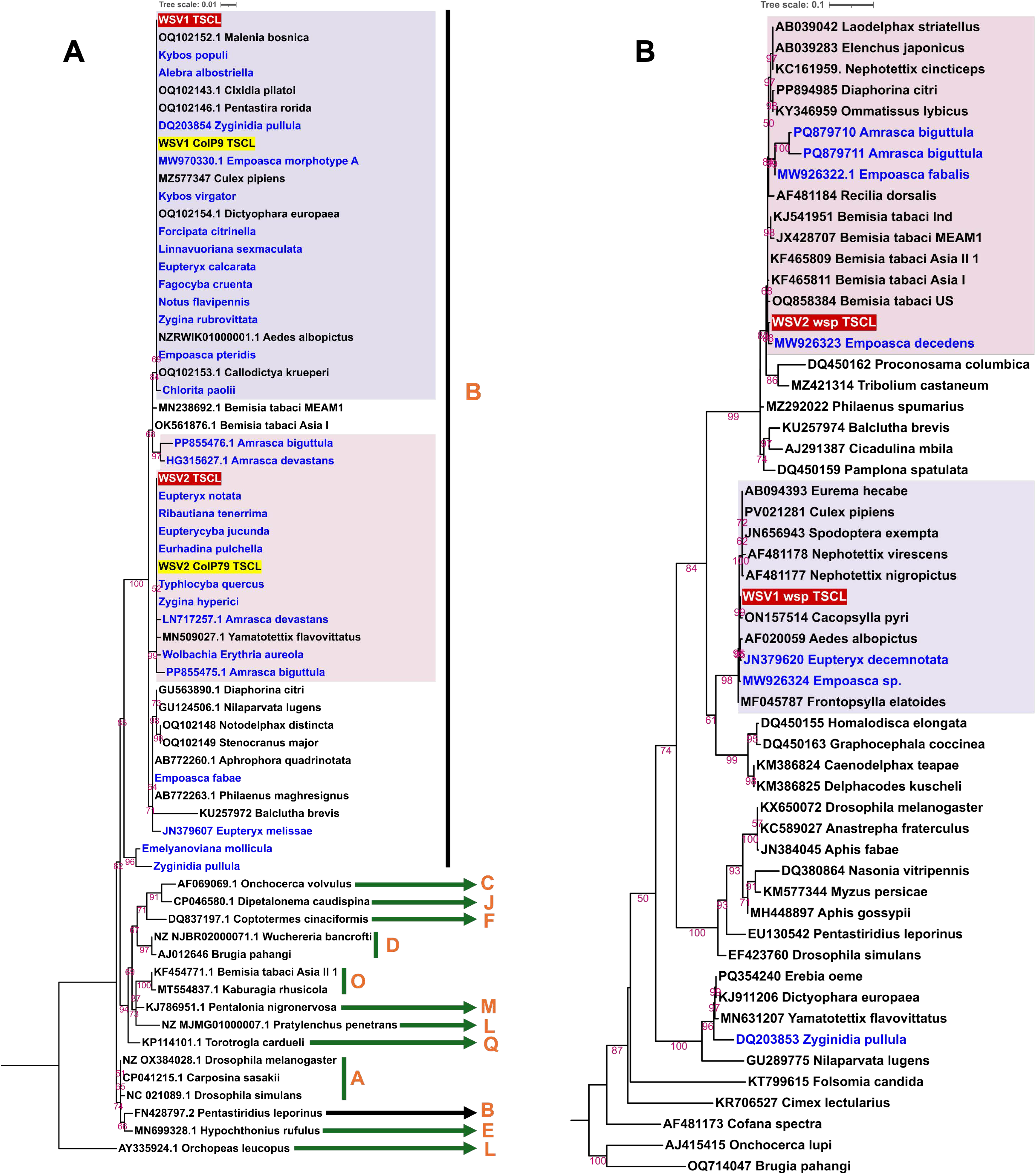
Phylogenetic relationship with other *Wolbachia* strains based on aligned 405 bp of 16S rDNA (A) and 461 bp of *Wsp* (B) nucleotide sequences. Nucleotide substitution models HKY+F+G4 and TPM3u+F+G4 were used for 16S rDNA and Wsp, respectively. *Wolbachia* strains from *Typhlocybinae* members (17) are indicated in blue and sequence variants identified by amplicon sequencing and PCR-RFLP in this study are highlighted in red and yellow, respectively.

### Localization of *Wolbachia* within TSCL

FISH analysis on wholemount TSCL adults localized *Wolbachia* signals primarily on the midgut region (Fig. 3A), whereas no signal could be detected from control insects without probe (Fig. S2A). This was confirmed by immuno-histochemistry on dissected midguts of TSCL using *Wolbachia* FtsZ polyclonal antibody showing localization in the conical midgut, midgut loop, and midgut posterior regions (Fig. 3 B, C) (N=4). No signal could be detected from a subset (N=5) of midguts (Fig. S2B), confirming the absence of *Wolbachia* in some individuals consistent with PCR results. No signal could be observed in control midguts stained only with the secondary antibody.

### Virus diversity in TSCL

BLASTX query of *de novo* assembled contigs from processed TSCL RNAseq reads against the non-redundant virus protein database yielded no detectable RNA viruses. Instead, 507 viral contigs with high similarity to dsDNA viruses belonging to the genus *Adintovirus* (*Eupolintoviridae*) were identified. From these, eight near full-length Adintovirus genomes (12.9-17.3 kb), designated as TSCL Adintovirus 1-8 were assembled (Fig.4). Reads of TSCL Adintovirus 7 was most abundant, followed by Adintovirus 4 and 6 (Table S3A). Annotation of the genome using BLASTP, CDD and HHpred algorithms revealed a characteristic genome organization of adintoviruses, comprising ORFs encoding DNA polymerase type B (Pol B), IS481 transposase, Integrase, Filament temperature-sensitive mutant K (FtsK), Hexon, Penton, GasderminX, Cupiennin, Phospholipase A2-like domain (PLA2X), Adenain, Malestrom (MAEL) and Phage T7 tail fiber proteins (Fig. 4). Pairwise comparison of PolB amino acid sequences of the TSCL adintoviruses revealed that the abundant TSCL Adintovirus 7 belonged to betadintoviruses sharing 10.9% and 39.3% (Table S3B) identity with reference sequences of *Mayetiola barley midge adintovirus* (*Alphaadintovirus*, <u>YP_010796961.1</u>) and *Terrapene box turtle adintovirus* (*Betaadintovirus*, <u>YP_010796957.1</u>), respectively. The other seven viruses shared higher identity (>36%) with alphadintoviruses (Table S3B). The presence of two abundant alphadintoviruses (TSCL adintovirus 4 and 6) and the betadintovirus (TSCL adintovirus 7) was validated by PCR amplification from DNA samples extracted from individual TSCL adults (N=17) using PolB and Hexon (major capsid protein) specific primers. All three Adintoviruses screened in this study were detected from 100% of tested samples.

**Fig. 4:**
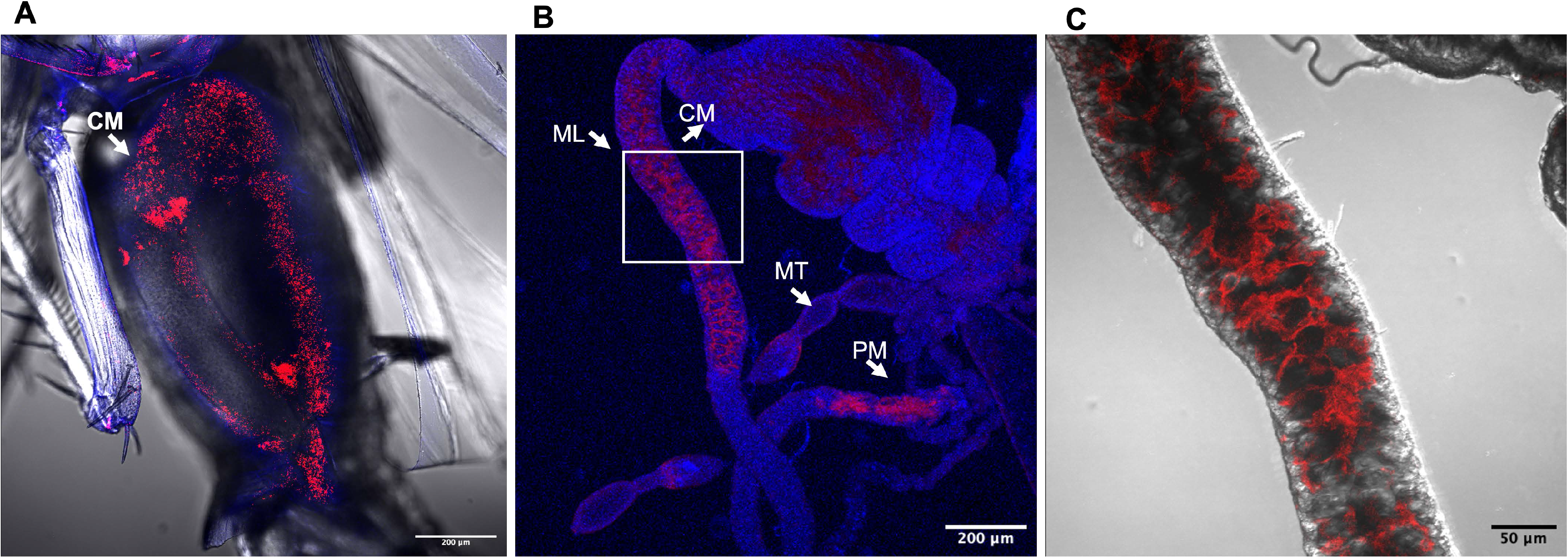
TSCL adult wholemount showing *Wolbachia* (red) and DAPI stained nuclei in midguts using FISH with cyanine3 labelled *Wolbachia* specific DNA probes (A), and immuno-localization of Wolbachia (red) in midguts using *Wolbachia* FtsZ polyclonal antibody (B) and magnified (20x) area under inset (C). CM, ML, PM and MT denote conical midgut, midgut loop, posterior midgut and Malpighian tubule, respectively.

**Fig. 5:**
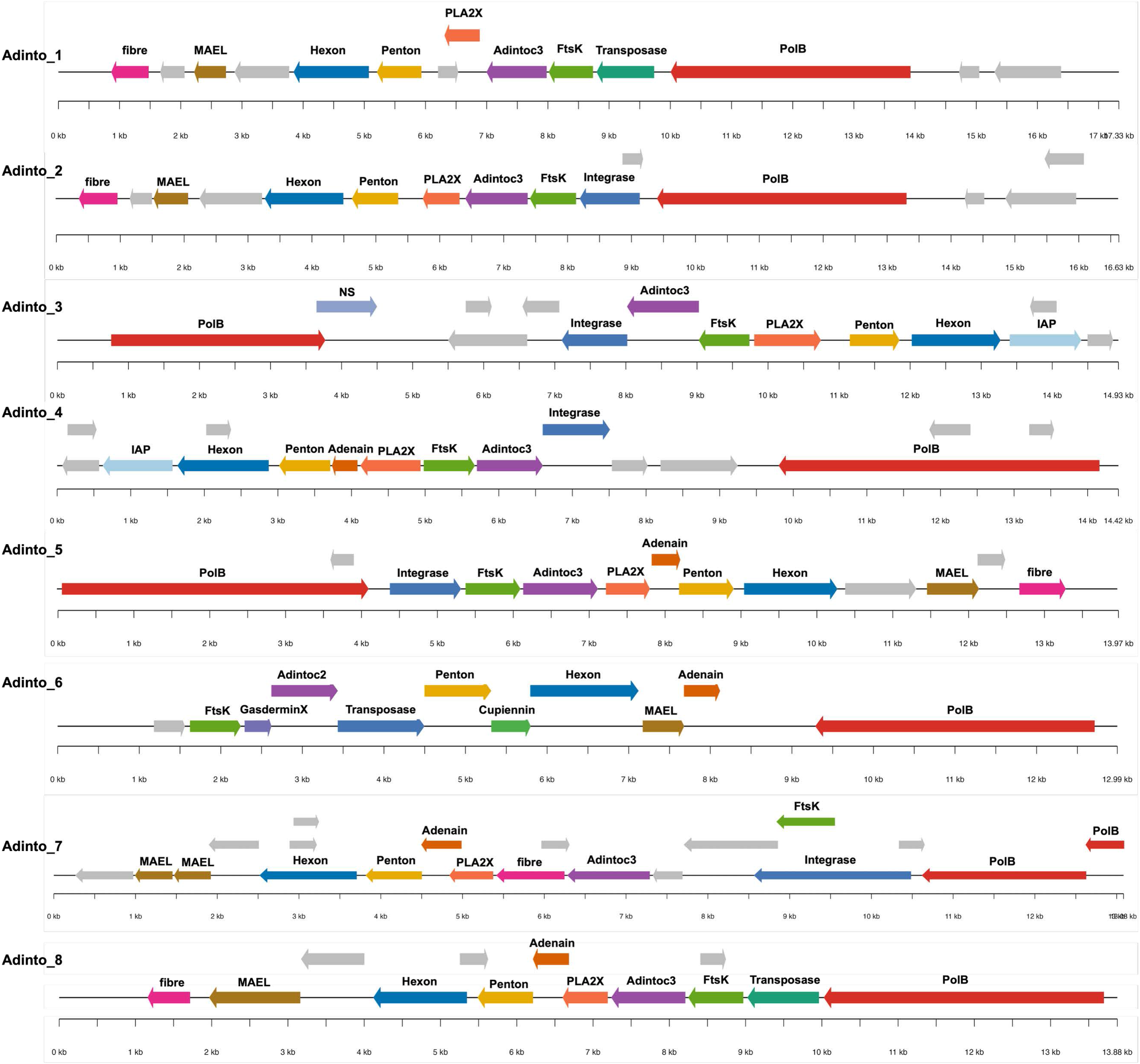
Genome organization of the full-length *Adintovirus* genomes (*Adintovirus* 1-8) identified in this study. Predicted open reading frames (ORFs) are represented with their orientation and encode DNA polymerase B (Pol B), IS481 family transposase, retrovirus-like integrase, double-jellyroll major capsid protein (hexons), single-jellyroll minor capsid protein (pentons) and Filament temperature-sensitive mutant K (FtsK), GasderminX, Cupiennin, Phospholipase A2-like domain (PLA2X), Adenain, Malestrom (MAEL), baculoviral inhibitor of apoptosis protein (IAP), AdintoC3, AdintoC2 and Phage T7 tail fiber proteins. The grey-colored boxes indicate ORFs that didn’t show clear hits in identity in searches using BLASTP, NCBI conserved domain database and HHpred.

**Fig. 6:**
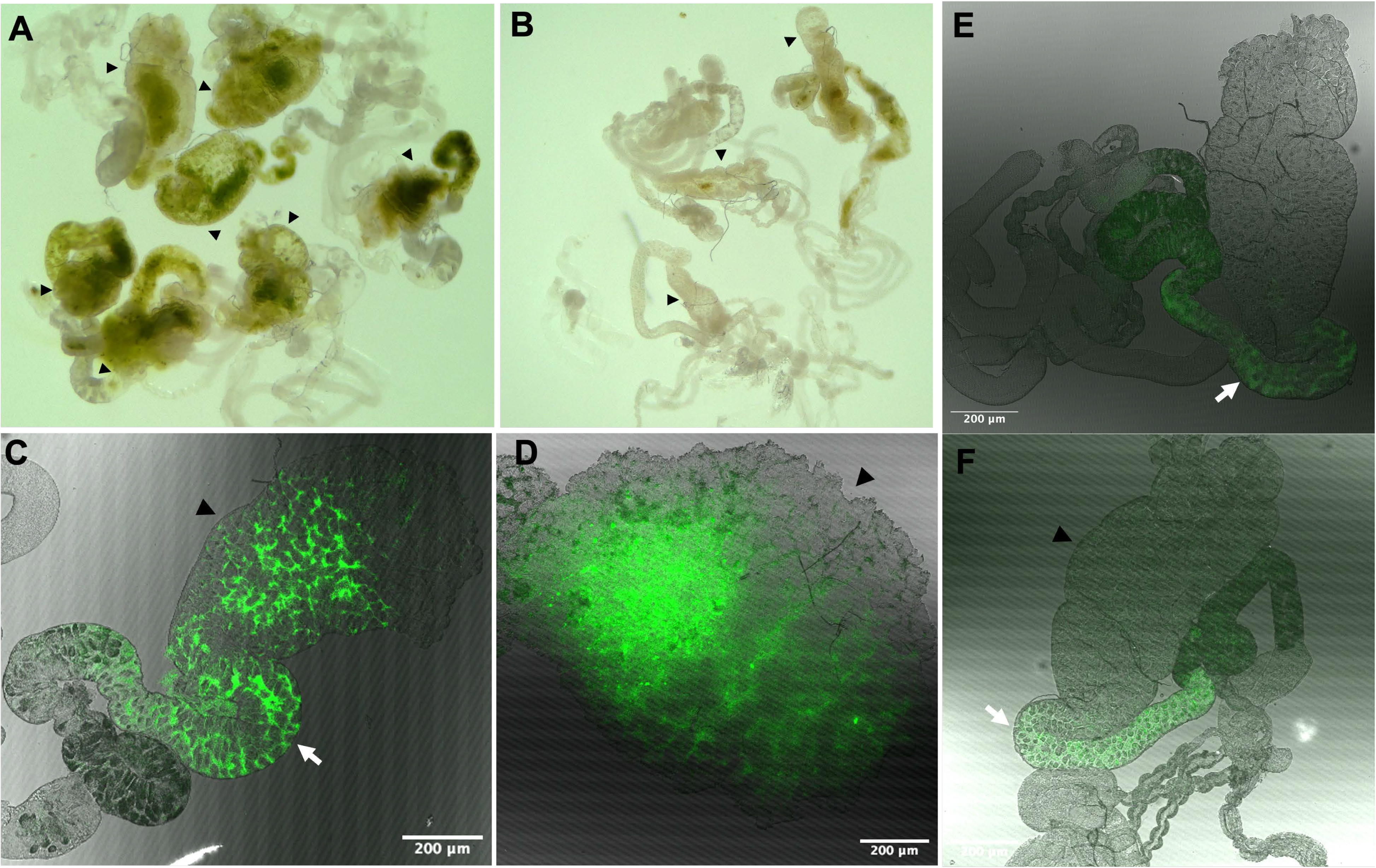
Midguts extracted from plant-fed adults appear more green with plant content (A) compared with control midguts (B) extracted from 15% sucrose diet fed adults with light microscopy. Autofluorescence detected using confocal microscopy in plant-fed (C, D) and diet-fed midguts (E, F) when excited with 555 nm wavelength. Black arrowheads indicating the conical midgut portions in plant fed treatments contained highest plant material and had higher autofluorescence.

### Fungal diversity

No symbiotic fungi could be identified from TSCL samples.

### Evidence of TSCL as mesophyll feeders

#### Autofluorescence of midguts

Midguts dissected from adults reared on cotton plants were green at the conical midgut region (Fig. 4A) compared with the control midguts (Fig. 4B) extracted from diet-fed adults. The presence of chlorophyll pigment was confirmed by autofluorescence detection from the conical midgut region (Fig. 4 C, D) of plant fed insects. Autofluorescence also was detected from the midgut loop region but not in the conical midgut region of control insects (Fig. 4 E, F) probably due to remnant chlorophyll pigments.

### Comparison of host plant chloroplast transcripts between pests of cotton

To further assess the mesophyll feeding nature of TSCL, normalized transcripts of cotton chloroplast genes identified from RNAseq data of TSCL, western flower thrips (epidermal feeder) and cotton aphid (phloem feeder) were compared. A total of 49 different chloroplast genes were identified from TSCL, compared with 23 and 4 genes identified from thrips and aphid, respectively (Fig. 7). Moreover, normalized transcript abundance of important chloroplast genes belonging to photosystem I, II, Rubisco, cytochrome b6, and ATP synthase complex was higher in TSCL compared with thrips and aphids (Fig. 7), indicating that plant mesophyll cells as the primary feeding niche for TSCL.

**Fig. 7:**
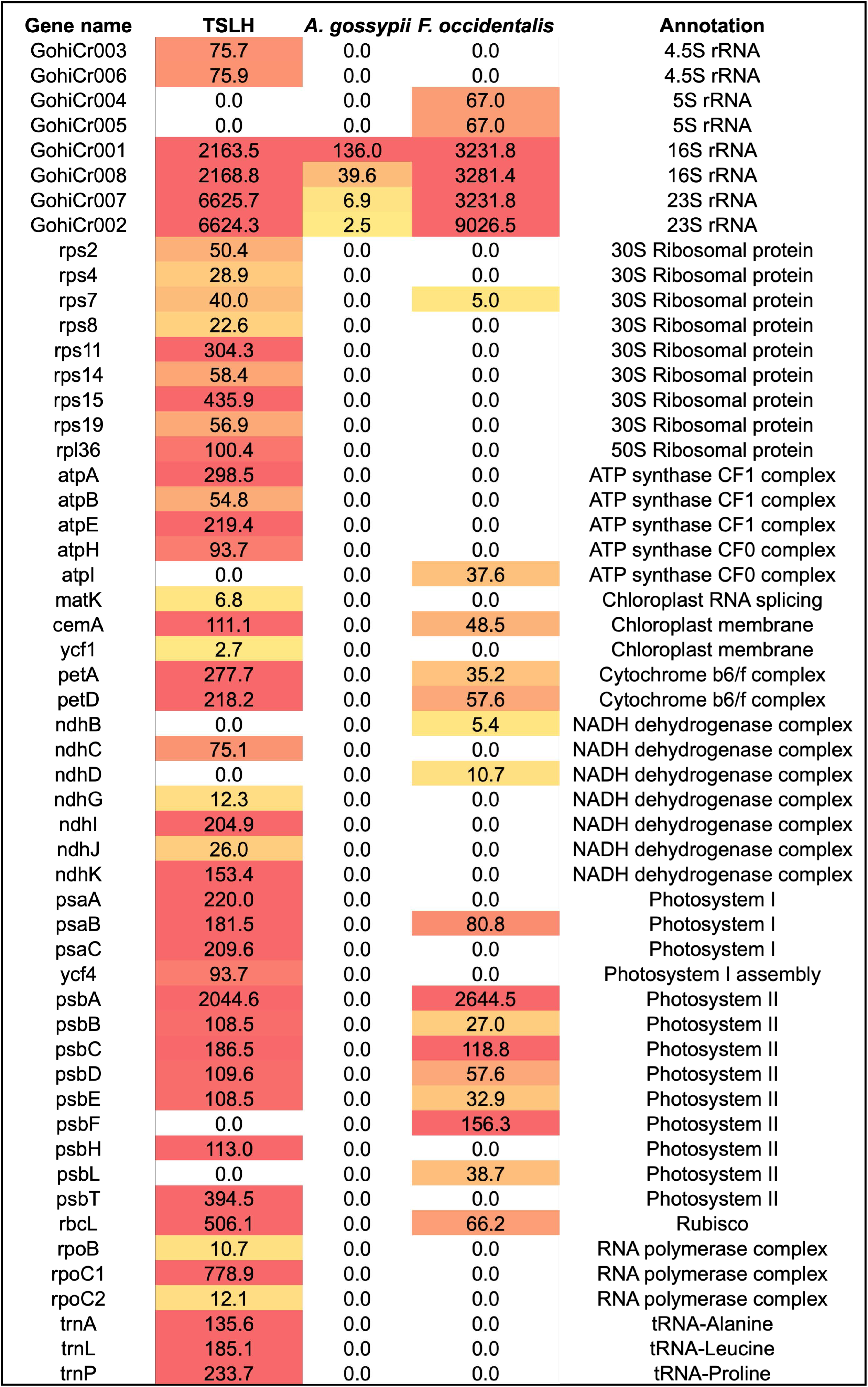
Comparison of abundance of normalized (transcripts per million) cotton chloroplast specific transcripts detected from RNAseq data generated from TSCL, cotton aphid (*Aphis gossypii*), and Western flower thrips (*Frankliniella occidentalis*). Comparative higher and lower abundance of normalized transcripts is indicated with red and yellow color shading.

#### EPG recordings

Waveforms recorded for TSCL probing were consistent but produced distinct patterns for adults (N=7) and nymphs (N=5), respectively (Fig. 8 A, B). Both indicated cell rupture style of probing characterized by rapid in and out stylet movement into the plant cells. The wave patterns in nymphs were more frequent but of lower amplitudes compared with that of adults (Fig. 9, 10). Waveforms recorded for adults included a regular sequence of spikes (A1 and A2) but at much lower frequency than the shorter spikes observed in case of nymphs (Fig. 9, 10). The A1a and A1b waveforms were regular and smooth at the tips but the latter had higher amplitudes. The A2 waveforms contained ruffles and were pointed at the tips. Brief span of potential drops was seen post A1 and A2 waveforms.

**Fig. 8:**
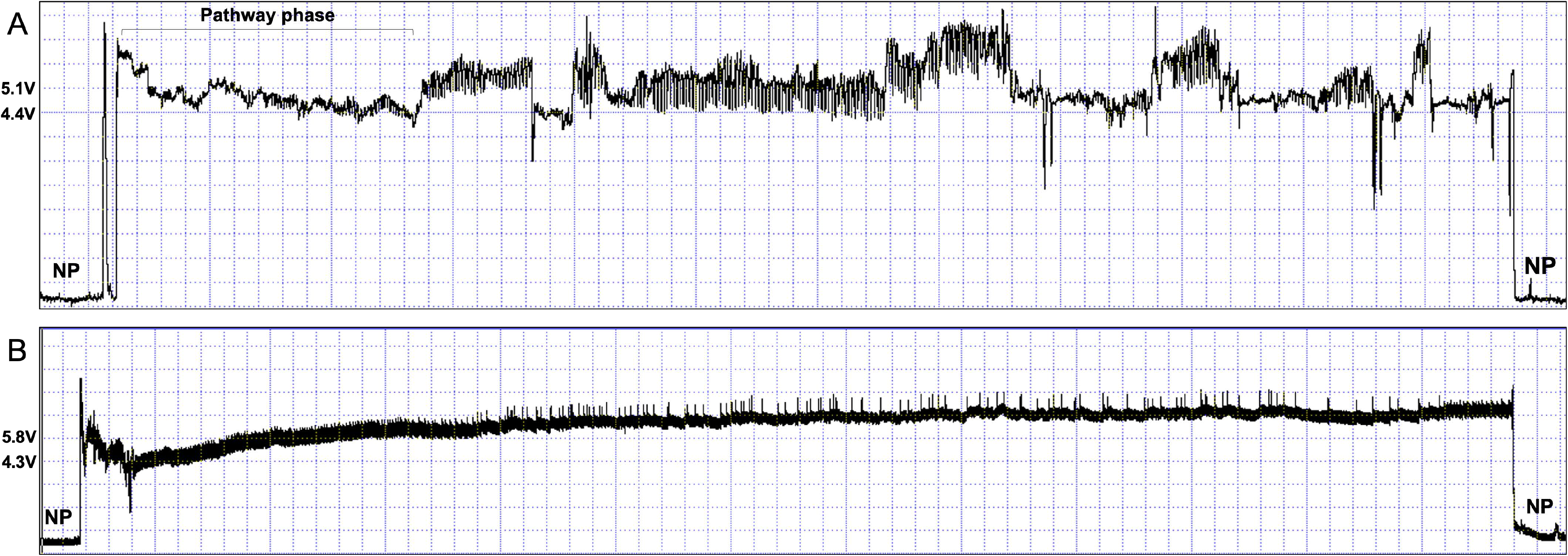
EPG pattern from start to end of a probe by TSCL adult (A) for a duration of 97 seconds at compression of 20 (1.67 seconds/division) and nymph (B) for 702 seconds at compression (11.2 seconds/division). NP denotes no probing. Vertical divisions indicate 0.71 and 1.56 V/division for adults and nymphs, respectively.

**Fig. 9:**
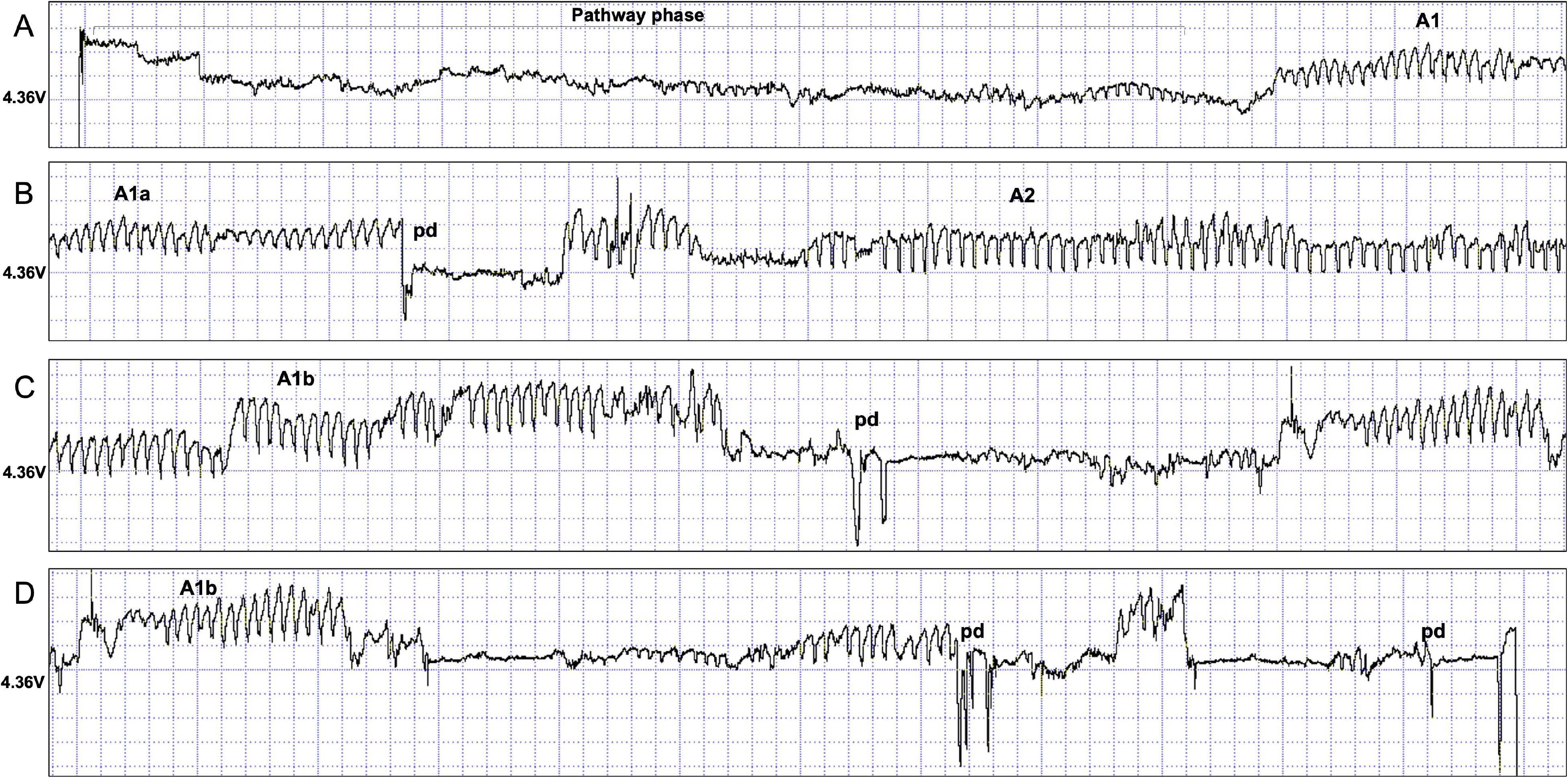
Illustrated characteristics of distinct waveforms identified from the adult probe at compression 5 (0.42 seconds/division) and 0.71 V/vertical division. pd denotes potential drop.

**Fig. 10:**
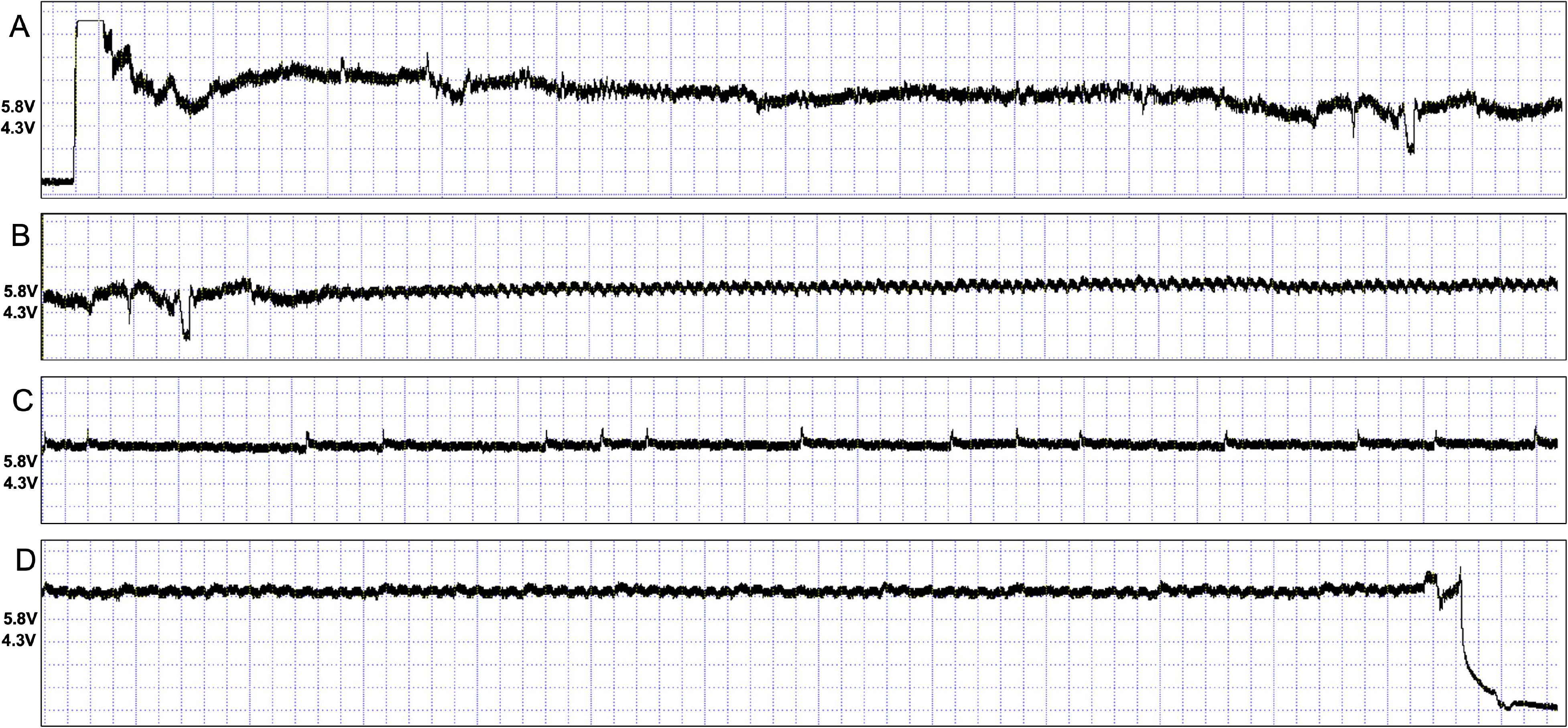
Illustrated characteristics of distinct waveforms identified from the nymph probe at compression 5 (0.42 seconds/division) and 1.56 V/vertical division.

### No evidence for TSCL transmitted pathogens causing hopperburn symptoms

Analysis of symptomatic plant tissues did not implicate a TSCL-transmitted pathogen as the cause of “hopperburn.” 16S rRNA sequencing revealed no known plant pathogenic bacteria, phytoplasmas, or *Spiroplasmas,* in symptomatic leaves. The bacterial communities of TSCL and field/lab collected host plant samples were significantly different and clustered separately in PCoA analysis (Fig. S3). Similarly, RNA-seq analysis of symptomatic plants did not identify any viruses associated with TSCL infestation, though Cotton leaf roll dwarf virus (CLRDV) was incidentally detected in field-collected cotton.

## Discussion

### Bacterial diversity

Leafhoppers belonging to *Typhlocybinae* sub-family exhibit low microbial diversity and in most cases are devoid of obligate/primary bacterial symbionts (17). Kobialka et al (17) identified *Wolbachia* as the predominant secondary bacterial symbiont within *Typhlocybinae* species. Our results showed that like other *Typhlocybinae* members, TSCL did not harbor any obligate bacterial or fungal symbionts, but *Wolbachia* was the predominant secondary bacterial symbiont. Reason for absence of obligate symbionts within this cicadellid sub-family remains unknown; however, their nutrient rich mesophyll diet could compensate for the lack of endosymbionts (17, 34). *Empoasca fabae*, another mesophyll feeding (33) *Typhlocybinae* pest of alfalfa, potato, and leguminous crops in the US, harbors *Sulcia* as a primary symbiont (36) alongside *Wolbachia* (53). Comparative studies of these two leafhopper species are imperative to understanding the impact of the absence of a primary symbiont on the host biology and feeding strategy.

This study revealed that *Wolbachia* is not fixed in the invading TSCL population screened in this study being detected in 60% of the individuals. Similar *Wolbachia* incidences (61.5 %) were previously reported from TSCL populations in Pakistan (54). Genetically diverse strains of *Wolbachia* belonging to B supergroup are harbored within *Typhlocybinae* members, including TSCL (17, 35, 36, 54, 55). Phylogenetic analysis in this study confirmed that two different strains of *Wolbachia* with varying abundance co-infect TSCL population screened in this study. Asymmetric densities of co-inhabiting *Wolbachia* variants are well-documented (56–58); however, factors accounting for the differential abundance of the two *Wolbachia* isolates in this study remains unknown. Notably, the lesser abundant WSV2 strain clustered together with *Wolbachia* strains previously identified from TSCL populations from Pakistan. Whereas the higher abundant WSV1 strain clustered differently. A recent study comparing phylogenetic relationships of global populations of TSCL using mitochondrial cytochrome oxidase markers revealed dominance of a single haplotype across native regions in South Asia and mainland USA (8). These findings reiterate the importance of studying microbial diversity together with insect genetic markers to understand ecological history of insects.

*Wolbachia* drives its spread by manipulating reproduction of its insect host (59), and evidence of *Wolbachia* indued cytoplasmic incompatibility and feminization is well documented in leaf/plant hoppers (35, 55, 60, 61). Our results show that *Wolbachia* reside within midguts of TSCL and is similar to previous reports of their midgut localization within *Typhlocybinae* members (17, 36). Midgut residing *Wolbachia* may supplement vitamins (62), induce immune response (63), or stabilize the midgut environment (64); however, its exact role in TSCL biology needs to be elucidated in future.

### Virus diversity

Virus diversity analysis of *Empoasca fabae* (65), *Matsumurasca onukii* (66) show predominance of RNA viruses belonging to *Iflaviridae*, *Solemoviridae*, *Lispiviridae*, *Phenuiviridae* and *Nidovirales*, compared with a lone DNA virus (*Parvoviridae*). Recently Iflaviruses (+ve, ssRNA) have been identified from TSCL samples from India (67) and China (<u>MZ367599.1</u>). Surprisingly, no RNA virus could be detected within the TSCL samples analyzed in this study, but eight diverse full length dsDNA genomes (12-17 kb) of Adintoviruses (TSCL Adintovirus 1-8) were identified from the RNAseq data. Adintoviruses were initially considered as polintons aka mavericks, which are large dsDNA transposons widespread among eukaryotic genomes and identified by presence of characteristic genes encoding protein-primed type B DNA polymerase (PolB) and retroviral-like integrase (68). However, recent characterization of double-jellyroll major capsid proteins (hexons), single-jellyroll minor capsid protein (pentons), and virus genome packaging ATPases (FtsK) (68) have them redefined as viruses belonging to newly proposed family, *Adintoviridae* (69). Adintoviruses are further classified as alphadintoviruses and betadintoviruses, based on the identity of the PolB sequences with reference sequences of PolB (pfam03175) in CDD database. Our results show that out of the eight adintoviruses detected from TSCL, seven belonged to alphadintovirus genera, whereas the most abundant (TSCL adintovirus 7) belonged to betadintovirus class. It is noteworthy that TSCL adintovirus 6 genome encodes characteristic beta-class accessory proteins such as GasderminX and Cupiennin together with an alpha-class PolB, suggesting possible horizontal gene transfer between alpha and beta class adintoviruses (69).

Adintoviruses can live a dual lifestyle of an endogenous virus element that can be induced to produce infectious virus particles (70). Interestingly, *Spodoptera* adintovirus genomes within Sf9 cells, a standard insect cell culture line, produced infectious virions that could infect High Five cells, –another standard insect cell line (69). Whether TSCL adintoviruses live as endogenous viruses integrated within the host genome or can form infectious particles remains to be resolved. Even if adintoviruses only remain as long-term heritable endogenized elements, mapping its genetic diversity within global TSCL populations can shed light on the ecological and evolutionary history of its insect host (71).

### Evidence for mesophyll feeding

This study demonstrates the mesophyll feeding nature of TSCL, using microscopic evidence of higher chlorophyll derived autofluorescence in midguts extracted from plant fed adults compared with control midguts extracted from artificial diet fed adults, and the presence of higher numbers of chloroplast derived RNA transcripts within RNAseq data of TSCL compared with phloem/epidermal feeders. In higher plants, mesophyll cells contain more and larger chloroplasts compared with epidermal or phloem parenchyma cells (72, 73), accounting for the higher abundance of chloroplast transcripts identified within mesophyll feeding TSLH compared with epidermal and phloem feeders, in this study.

Mesophyll feeding *Typhlocybinae* members use rapid ‘in and out’ movement of its stylet to rupture plant cells and ingest cell contents (74, 75). This unique stylet movement along with secreted salivary effectors, induces plant wound responses leading to hopperburn symptoms (75). Electropenetrography (EPG) patterns for TSCL adults and nymphs in this study clearly show similar stylet movement indicative of cell rupture feeding. Interestingly, distinct wave patterns were observed between adult and nymphs, with highly rapid stylet movements of low amplitude seen in the case of nymphs. TSCL nymphs are believed to cause more prominent symptoms compared with adults (1, 2) and this could be due the differences noted here in the feeding pattern. Hunter et al. (33) describe three different waveforms I_a,_ I_b_ and I_c_ of the mesophyll feeding *Empoasca fabae*, through EPG, wherein high amplitude peaks (I_a_) indicated lacerating stylet movement, I_b_ corresponded to single spongy mesophyll cell puncture and feeding, and I_c_ indicated ingestion activity. Although the exact plant tissue associated with ingestion (I_c_) feeding was not determined, duration of I_c_ events has been implicated with mesophyll (< 2 min) or phloem feeding (>2 min) activity of the potato leaf hopper. The waveforms A1 and A2 described in this study are very similar to the Ia and Ib waveforms described previously (33, 74, 76). However, no clear Ic like waveform was identified from EPG recordings in this study. It is important to note that phloem restricted plant pathogens such as phytoplasmas (77, 78) and mastreviruses (79) have been detected from TSCL, suggesting that apart from the mesophyll, TSCL also feeds on phloem tissues. Thorough characterization of EPG waveforms described in this study together with microscopy is needed to decipher extent of mesophyll and phloem feeding by TSCL. Nevertheless, TSCL feeding induced symptoms (Fig. S4) are likely a cumulative effect of its cell rupture feeding style and secreted salivary effectors. Experimental evidence confirming systemic effect of TSCL secreted saliva and functional characterization of salivary proteins is essential to understanding mechanisms associated with hopperburn symptoms.

### Genbank accession numbers

Nucleotide sequences generated in this study have been deposited in NCBI Genbank under accession numbers PZ545711-17 (16S rDNA), PZ547707-12 (Wsp) and PZ547713 (FtsZ). Raw sequence reads generated in this study for amplicon sequencing of bacterial 16S rDNA of TSCL and host plants; fungal ITS; RNAseq of TSCL, host plants and thrips have been submitted to NCBI under the bioproject PRJNA1490287.

## Supplementary figure legends

**Fig. S1** Phylogenetic relationship with other *Wolbachia* strains based on aligned 411 bp *FtsZ* (cell division protein) nucleotide sequences using TPM3u+F+G4 nucleotide substitution model.

**Fig. S2:** Wholemount control without probe for detection of *Wolbachia* using FISH (A). Midguts showing no signal for *Wolbachia* using FtsZ polyclonal antibody and Alexa Fluor^TM^ 488 labelled anti-rabbit secondary antibody (B).

**Fig. S3:** Principal co-ordinate analysis (PCoA) of Bray-Curtis dissimilarities showing absence of common bacterial genera within TSCL and host plants and proves no possible involvement of TSCL transmissible bacteria as the possible causal agent of hopperburn symptoms.

**Fig. S4:** Hopper burn symptoms in cotton plants reared with laboratory population of TSCL (A) and mesophyll clearing on adaxial and abaxial sides of leaves (B) attached with clip cages with TSCL adults (N=3) after 4 days of feeding.

## Supplementary table legends

**Table S1:** Primers used in this study for PCR amplication of 16S rDNA, Wsp (Wolbachia surface protein), FtsZ (cell division protein), adintovirus hexon, and polymerase B (PolB) genes **Table S2:** Abundance of bacterial genera identified by amplicon sequencing of V3-V4 (A) and full length 16S rDNA (B), and *Wolbachia* sequence variants obtained from 16s full length 16S rDNA.

**Table S3**: Mean read depth and abundance of Adintovirus contigs identified in this study (A). Annotation of DNA polymerase B (Pol B) genes against assigned Pfam families and pairwise identity RefSeq sequences of *Mayetiola barley midge adintovirus* (*alphadintovirus,* YP_010796957.1) and *Terrapene box turtle adintovirus* (Betadintovirus, YP_010796961.1) (B).

