## Supplementary figures and images for "Microbial diversity and insights into feeding behavior of the two-spot cotton leaf hopper (*Amrasca biguttula*)"

### Figure S1

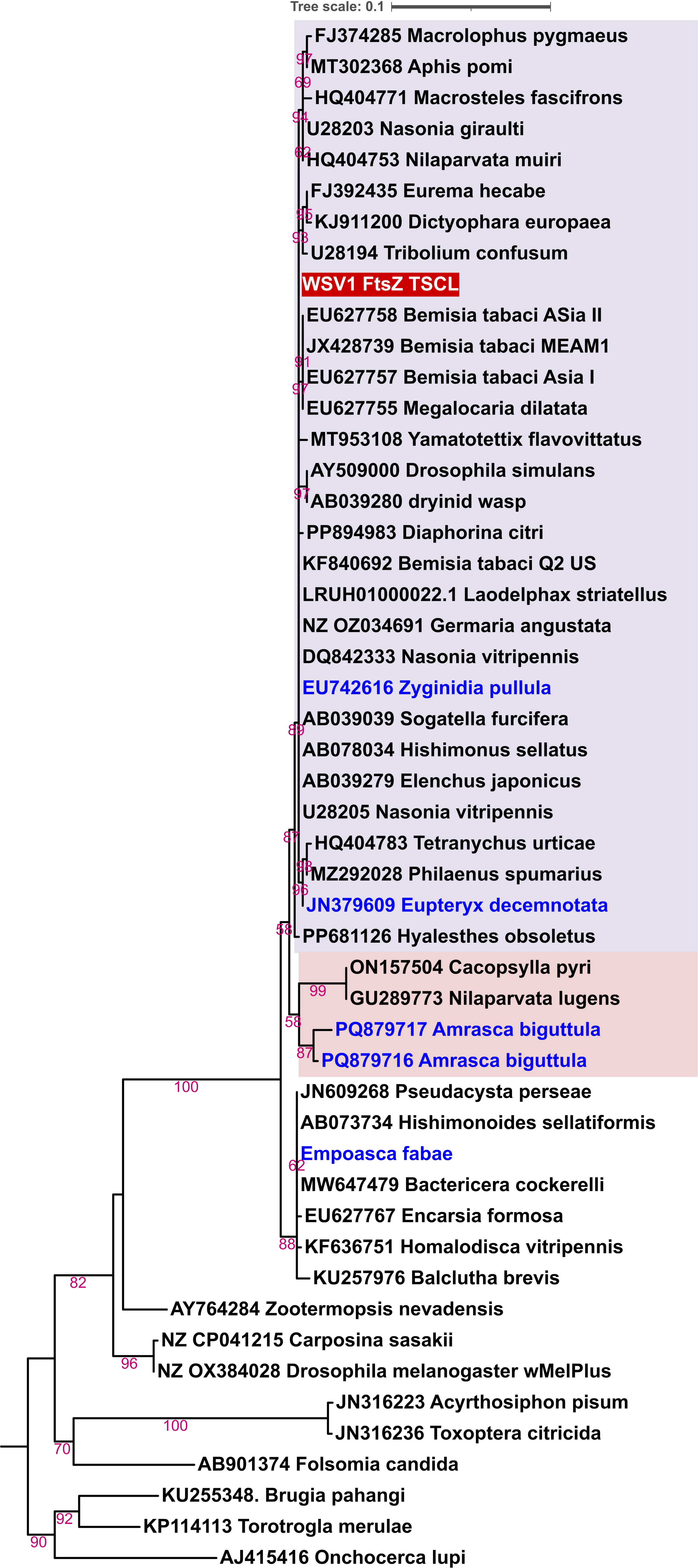

### Figure S2

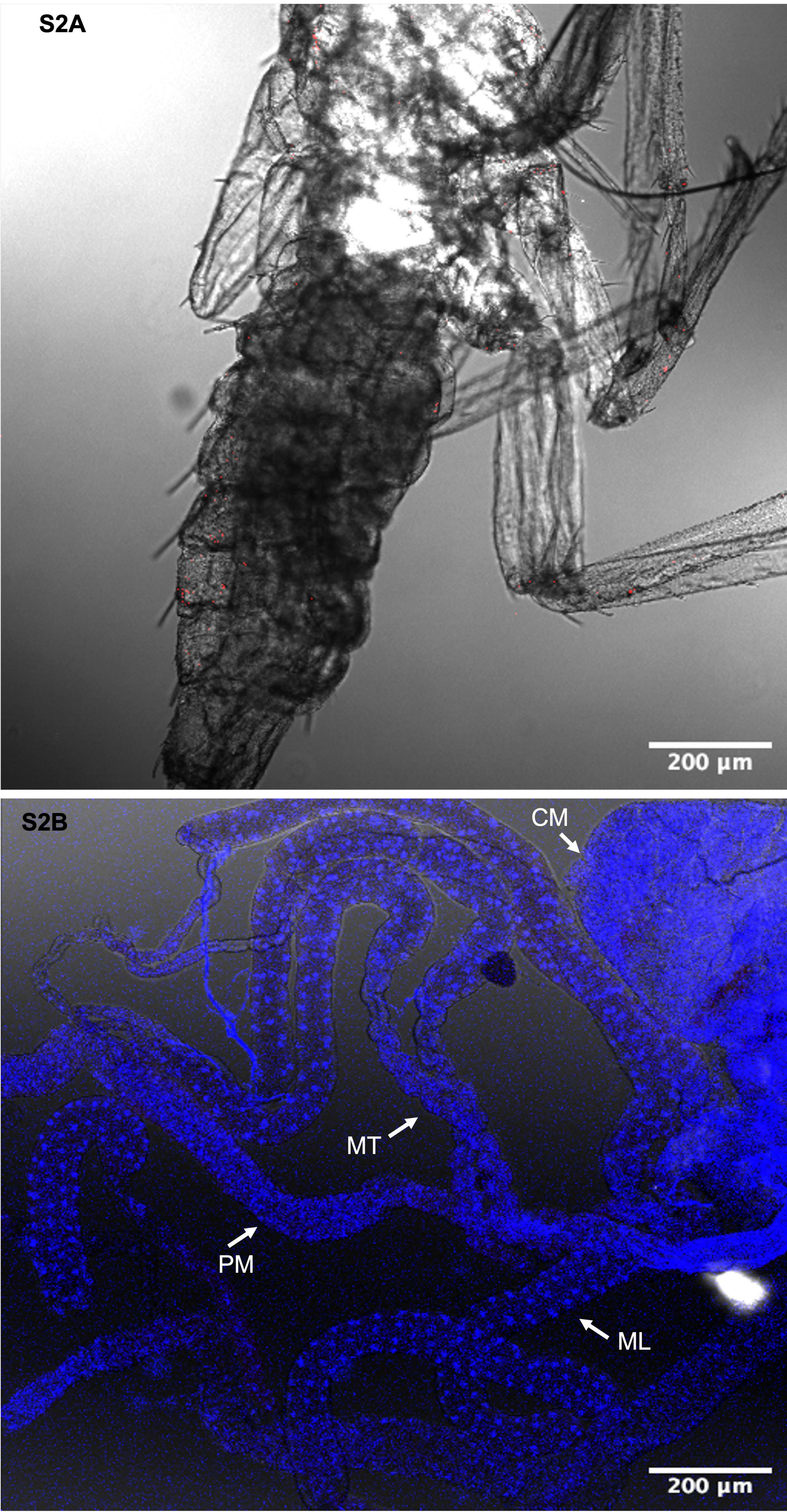

### Figure S3

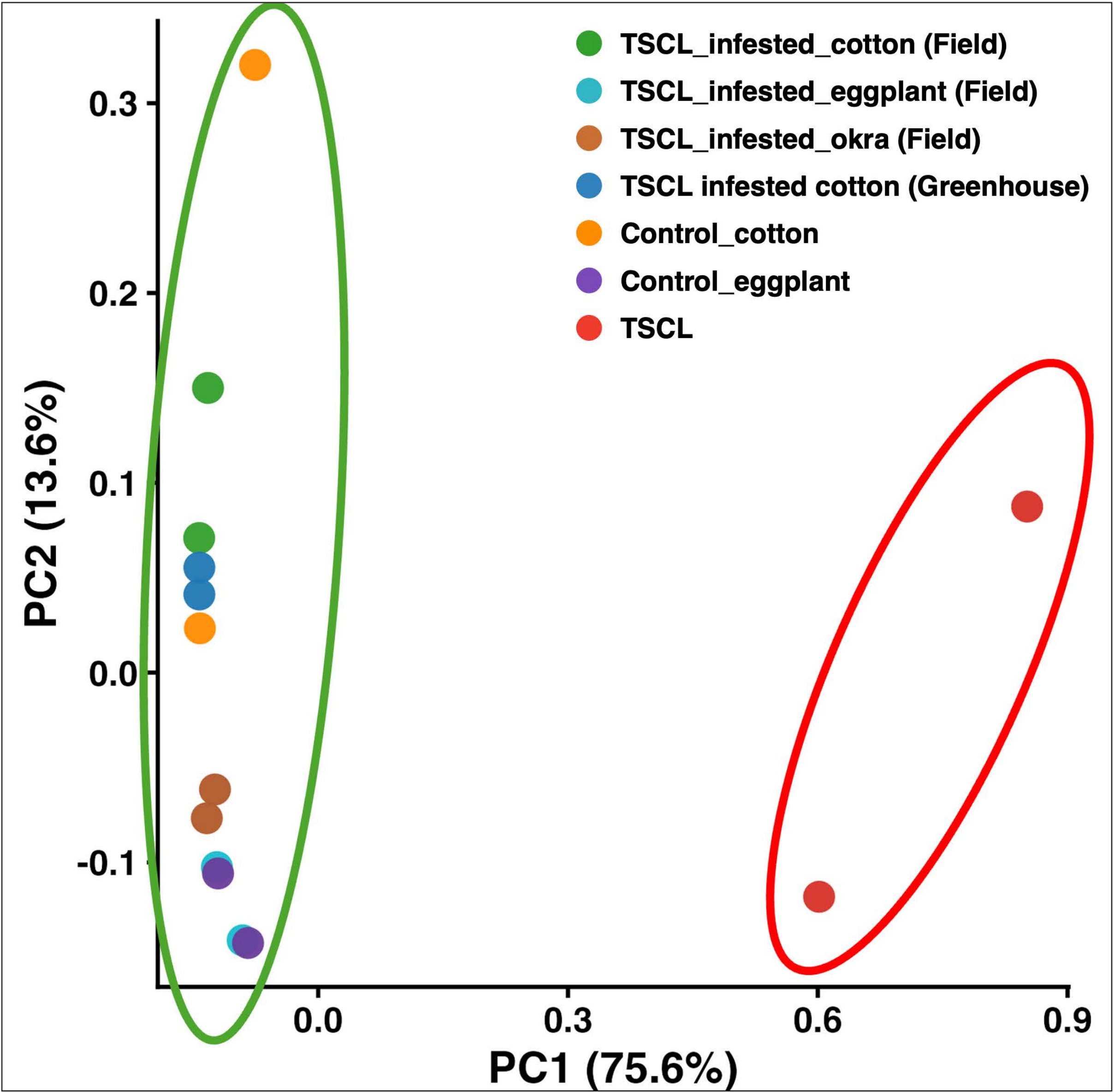

### Figure S4

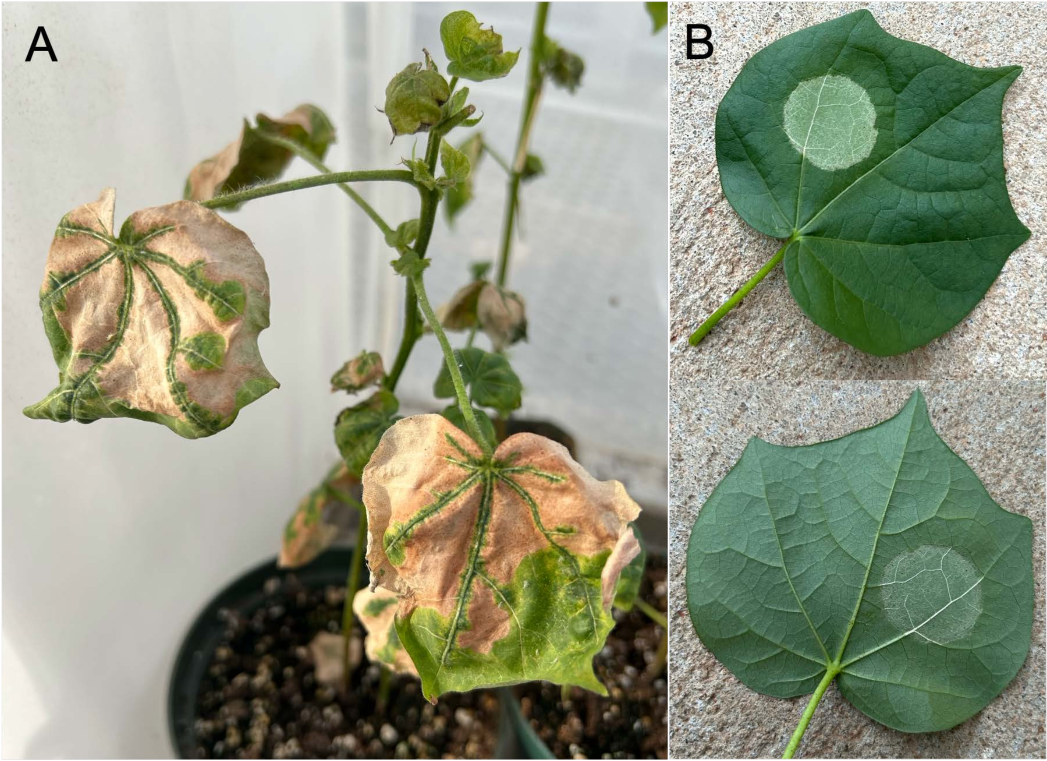
